# The maize gene *maternal derepression of r1* (*mdr1*) encodes a DNA glycosylase that demethylates DNA and reduces siRNA expression in endosperm

**DOI:** 10.1101/2021.10.04.463062

**Authors:** Jonathan I. Gent, Kaitlin M. Higgins, Kyle W. Swentowsky, Fang-Fang Fu, Yibing Zeng, Dong won Kim, R. Kelly Dawe, Nathan M. Springer, Sarah N. Anderson

## Abstract

Demethylation of transposons can activate expression of nearby genes and cause imprinted gene expression in endosperm, and it is hypothesized to lead to expression of transposon siRNAs that reinforce silencing in the next generation through transfer either into egg or embryo. Here we describe *maternal derepression of r1* (*mdr1*), which encodes a DNA glycosylase with homology to Arabidopsis *DEMETER* and which is partially responsible for demethylation of thousands of regions in endosperm. Instead of promoting siRNA expression in endosperm, MDR1 activity inhibits it. Methylation of most repetitive DNA elements in endosperm is not significantly affected by MDR1, with an exception of Helitrons. While maternally-expressed imprinted genes preferentially overlap with MDR1 demethylated regions, the majority of genes that overlap demethylated regions are not imprinted. Double mutant megagametophytes lacking both MDR1 and its close homolog DNG102 result in early seed failure, and double mutant microgametophytes fail pre-fertilization. These data establish DNA demethylation by glycosylases as essential in maize endosperm and pollen and suggest that neither transposon repression nor genomic imprinting are its main function in endosperm.

## INTRODUCTION

In 1970, Jerry Kermicle reported that the *red-color* (*R*) gene required for pigmentation in maize endosperm was preferentially expressed from maternal alleles (Kermicle 1970). Since then, genomic imprinting has been reported for hundreds of genes in maize, other plants, and animals as well [reviewed in (Batista and Köhler 2020)]. Research primarily on *Arabidopsis thaliana* endosperm has revealed that reduction of DNA methylation of the maternal genome can differentiate maternal and paternal alleles. In Arabidopsis, a key enzyme is a DNA glycosylase called DEMETER, which cleaves the base-sugar bond in 5-methylcytosine leading to demethylation. It is primarily active in the central cell of the megagametophyte, which gives rise to the endosperm upon fertilization, rather than in the endosperm itself (Choi et al. 2002; Gehring et al. 2006; Park et al. 2017). In the pollen vegetative nucleus, where DEMETER is also highly active, many transposons are demethylated and expressed (Schoft et al. 2011; He et al. 2019). DNA methylation not only represses transposons, but can also regulate developmental gene expression, presumably through inhibiting or promoting transcription factor binding (O’Malley et al. 2016; Batista et al. 2019; Borg et al. 2021; Khouider et al. 2021). Absence of maternal DEMETER causes 100% seed inviability (Choi et al. 2002). Pollen that lack DEMETER give rise to apparently normal seeds, but have reduced transmission frequency relative to wild-type and have pollen tube defects, severity dependent on the ecotype (Xiao et al. 2003; Schoft et al. 2011; Khouider et al. 2021). In rice, the DNA glycosylase ROS1a/OsDNG702 is also strictly required for seed development and has a function in pollen fertility, but its importance in pollen may be dependent on the cultivar (Ono et al. 2012; Zhou et al. 2021). The Arabidopsis genome also encodes three other glycosylases, which along with *demeter* mutants, produce subtle phenotypes in a variety of plant tissues (Pillot et al. 2010; Yamamuro et al. 2014; Schumann et al. 2017; Schumann et al. 2019; Khouider et al. 2021; Kim et al. 2021). Endosperm derived from central cells that lack DEMETER have excessive DNA methylation at thousands of small regions (Ibarra et al. 2012; Park et al. 2016). Unifying features of these regions have not been identified, but they exhibit a preference for short transposons in genic areas and for the ends of long transposons. What guides DEMETER to these regions is unknown, but involves FACT, a versatile ATP-dependent chromatin remodeling complex (Frost et al. 2018). DEMETER depends on FACT for activity at about half of its ∼10,000 identified target regions, which are distinguished by being more heterochromatic than the other half. The N-terminal half of DEMETER is likely important for targeting, as truncation of the protein decreases its activity at normal targets and leads to increased activity in gene bodies (Zhang et al. 2019).

In addition to maternal demethylation by DEMETER in specific regions, there is a pronounced and dynamic global reduction in DNA methylation of both parental genomes in endosperm relative to typical vegetative tissues in Arabidopsis (Hsieh et al. 2009; Ibarra et al. 2012; Moreno-Romero et al. 2016). A similar phenomenon has been observed in other plants, e.g., rice and in *Brassica rapa* (Park et al. 2016; Chakraborty et al. 2021). This global reduction in methylation is not caused by DNA glycosylase activity, but instead by passive demethylation through DNA replication and reduced expression of multiple components of RNA-directed DNA methylation (RdDM) and the replication-coupled methyltransferase MET1 (Hsieh et al. 2011; Jullien et al. 2012; Belmonte et al. 2013; Kawakatsu et al. 2017). RdDM activity apparently increases later in endosperm development as evidenced by an increase in DNA methylation in all sequence contexts after cellularization (Moreno-Romero et al. 2016). Polycomb Repressive Complex2 (PRC2) promotes non-CG methylation in endosperm but is itself inhibited by DEMETER. Thus, while the direct effect of DEMETER is to reduce CG and non-CG methylation, it has a genome-wide effect of increasing non-CG methylation in endosperm (Hsieh et al. 2009; Ibarra et al. 2012). PRC2 also functions in imprinting of many genes independently of DEMETER [reviewed in (Batista and Köhler 2020)]. How much of endosperm demethylation is determined by the central cell is unclear. The central cell is unusual and distinct from endosperm in its lack of H3K9me2 and visible chromocenters (Garcia-Aguilar et al. 2010; Pillot et al. 2010; Yelagandula et al. 2014). It has apparently normal methylation outside of DEMETER target regions, however (Park et al. 2016).

Maternal demethylation of specific transposons can activate nearby gene expression [reviewed in (Anderson and Springer 2018; Batista and Köhler 2020)]. Studies in multiple species have revealed a high level of variation in which genes are imprinted, including within-species variation, and in some cases imprinting is associated with presence of transposons near genes (Gehring et al. 2009; Waters et al. 2013; Pignatta et al. 2014; Hatorangan et al. 2016; Pignatta et al. 2018; Rodrigues et al. 2021). Demethylation of transposons in the central cell is hypothesized to reinforce transposon silencing in the embryo (Ibarra et al. 2012; Bouyer et al. 2017). In this model, transposon activation produces siRNAs in central cell or endosperm that move into the egg cell or embryo. Likewise, DEMETER activity in the pollen vegetative cell could silence transposons in the sperm cells. (Calarco et al. 2012; Ibarra et al. 2012; Martínez et al. 2016). Methylation patterns in rice *ros1a* mutant pollen also support this hypothesis, where regions that are gain methylation in the mutant pollen vegetative cell lose methylation in the mutant sperm cell (Kim et al. 2019).

While work on gene regulation and chromatin in Arabidopsis endosperm has provided many insights into maize as well, maize is different than Arabidopsis in several ways. Maize lacks a CMT2-type chromomethyltransferase, causing CHH methylation (mCHH, where H is A, T, or C) to be largely depleted from heterochromatin and instead associated specifically with RdDM (Zemach et al. 2013; Gent et al. 2014). It has a much larger set of transposons, including ones in and near genes and many that are currently active (McCarty et al. 2005; Springer et al. 2018; Dooner et al. 2019). Its endosperm persists past fertilization and has a distinct cellular morphology (Becraft and Gutierrez-Marcos 2012). Unlike both Arabidopsis and rice endosperm, maize endosperm undergoes little demethylation genome-wide (Zhang et al. 2014; Fu et al. 2018). Kermicle’s demonstration that certain *R* haplotypes are activated specifically from the maternal genome in endosperm suggests that similar imprinting mechanisms function in maize as in rice and Arabidopsis. The phenomenon Kermicle studied is known as R mottling because of the pigmentation pattern. Maternal *R* is uniformly active and yields solidly pigmented endosperm. When maternal *R* is absent, non-uniform and weak expression of paternal *R* causes a mottling pattern. Kermicle also reported that mottling can occur in the presence of maternal *R* in a mutant he called *mdr* (for *maternal derepression of R*) (Kermicle 1995) (Figure 1A). He mapped the mutation to the long arm of chromosome 4, found it to be homozygous fertile, and demonstrated that maternal *mdr* activity is required and sufficient for uniform *R* activation in endosperm (Kermicle 1995). In this study, we mapped *mdr* to a single gene that encodes one of two similar glycosylases with endosperm expression in maize and demonstrated that it is required for full demethylation of thousands of diverse regions in endosperm.

**Figure 1.**
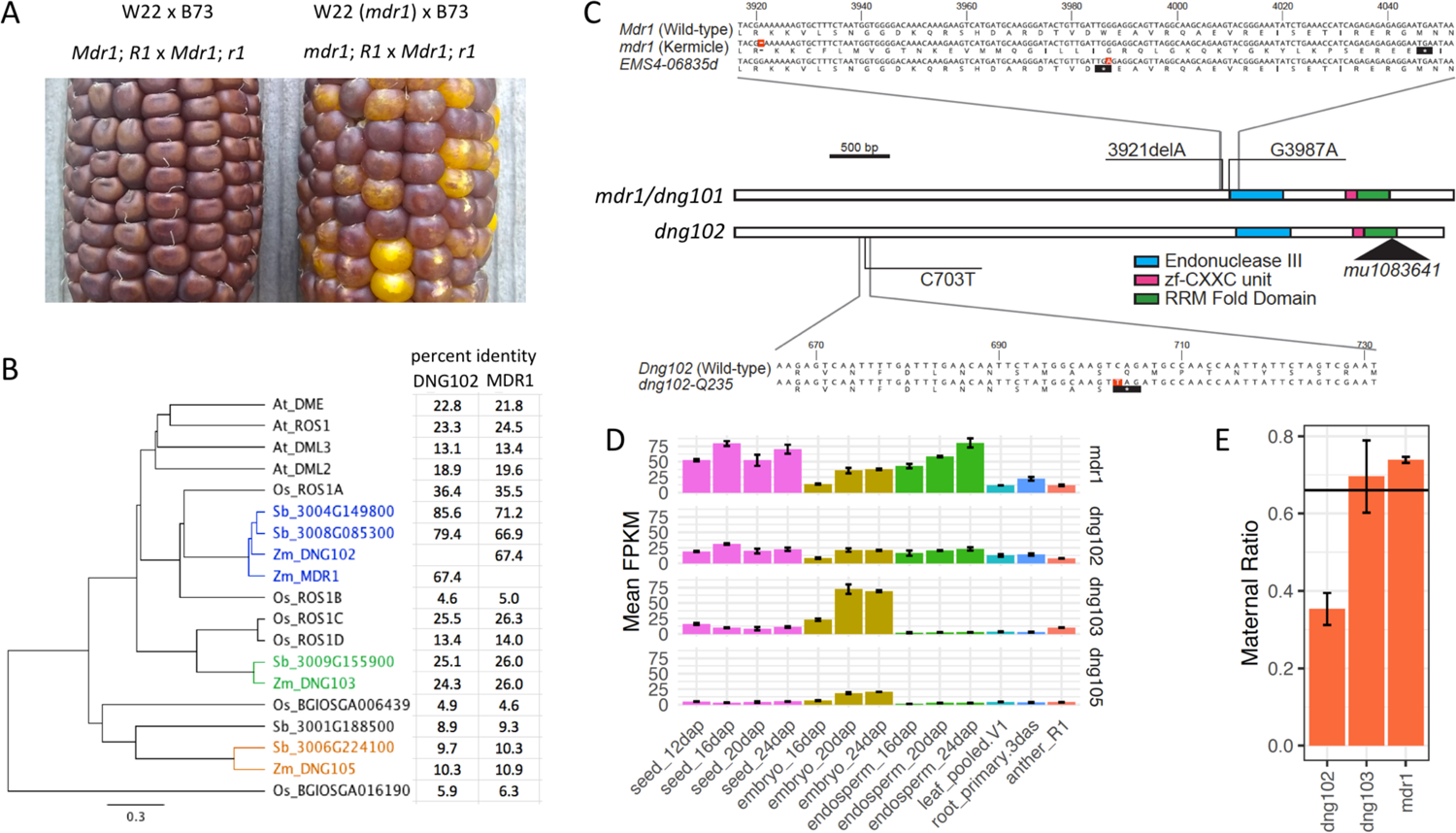
**A**, Kermicle’s mutant *mdr1* allele produces a maternal R mottling phenotype. In crosses where functional *R1* is inherited maternally only, maternal *Mdr1* is required for full *R1* expression. W22 (*mdr1*) is the *mdr1* mutant in a full-color W22 background. All listed alleles are homozygous. **B**, Protein tree of DNA glycosylases. The three types of *Zea mays* DNA glycosylases and *Sorghum bicolor* homologs are in separate colors, *Oryza sativa* and *Arabidopsis thaliana* in black. Percent identities are based on whole protein alignments. **C**, Sequence and structure of *mdr1* and *dng102* alleles. Nucleotides highlighted in red indicate positions of mutations. Asterisks in black background indicate positions of stop codons. Black triangle indicates position of Mu insertion. **D,** Expression of maize DNA glycosylases across tissues (Stelpflug et al. 2016). Error bars are standard deviations from the biological replicates. **E,** Maternal ratio of expression of DNA glycosylases in endosperm. Height of columns indicates proportion of maternal to paternal SNPs (Anderson et al. 2021). Horizontal line indicates expected ratio of 0.67 for non-imprinted genes. Error bars are standard deviations of the biological replicates. *dng105* is excluded because of lack of read coverage of informative SNPs.

## RESULTS

### The *maternal derepression of r1* (*mdr1*) gene encodes a DNA glycosylase

Consistent with current nomenclature in which the original *red-color* gene *R* was changed to *r1*, we refer to *mdr* as *mdr1* (https://www.maizegdb.org/nomenclature). To map *mdr1*, we took advantage of the fact that *mdr1* originated spontaneously in a W22 inbred stock (Kermicle, personal communication). We crossed *mdr1*/*Mdr1* heterozygotes as females with *mdr1* homozygotes as males to produce 50% homozygous and 50% heterozygous *mdr1* (Supplemental Figure S1A). After DNA extraction in bulk from ∼500 plants grown from mottled kernels, we Illumina sequenced to identify variants associated with mottling. For comparison, we also sequenced five individuals of the W22 stock that served as the source for a wild-type *Mdr1* allele. The presence of other spontaneous mutations that differentiate Kermicle’s W22 from the reference genome W22 made it possible to identify a region of approximately 20 MB on the distal tip of the long arm of chromosome 4 (4L) that had non-W22 alleles at nearly 100% frequency in mottled kernels. (Supplemental Figure S1B). Several other regions of the genome had unexpectedly abundant non-W22 alleles, which revealed where Kermicle’s W22 differed in ancestry from the reference W22 (Supplemental Figure S1C). Importantly, 4L was not among these regions, which greatly limited the number of candidate mutations. In fact, within the 20-Mb we identified only 45 SNPs (or short indels), and only one of them was in an exon. This was a deletion of a single A within a string of seven A’s, which created a frameshift mutation in *dng101* (*DNA glycosylase 101)* upstream of the predicted Endonuclease III and RRM-fold domains that are typical of DEMETER and ROS family glycosylases (Figure 1B, C) (Since *dng101* is encoded on the negative strand, this is a deletion of a single T in the genome sequence, at position 229,617,698). To confirm that the mutation in *dng101* caused the maternal R mottling phenotype, we also obtained an EMS allele (EMS4-06835d) from the Lu et al (2018) Gene-Indexed Mutations in Maize collection (Lu et al. 2018), which caused a G to A nucleotide change in the same exon (at position 229,617,638 in the genome), producing a premature stop codon. After backcrossing this allele four times into W22 to create an *EMS4-06835d* heterozygote in a W22 background, we tested for a mottling phenotype by crossing the heterozygote as the female parent with the wild-type *Mdr1* homozygous B73 inbred stock as male. Genotyping of *EMS4-06835d* in plants grown from these kernels revealed 20 of 20 plants from the strongest mottling kernels inherited the mutation while only 9 of 28 from solid kernels did (two tailed P-value < .0001, Fishers Exact test). These results provide independent evidence that mutations in *dng101* cause maternal R mottling. Thus, we refer to *dng101* as *mdr1*. Unless otherwise indicated, we use the Kermicle allele for all subsequent *mdr1* mutant experiments.

### A functional *Mdr1* allele or a functional *Dng102* allele is required for male and female fertility

Three additional genes in maize encode DEMETER-like Endonuclease III and RRM-fold domains, *dng102*, *dng103*, and *dng105*. See Supplemental Table S1 for transcript IDs for each gene. These make up three distinct subtypes, with *mdr1* and *dng102* together making up one subtype (67% amino acid sequence identity with each other) (Figure 1B, C). Each subtype has at least one homolog in sorghum. While sorghum also has two genes of the *mdr1*/*dng102* subtype, one of them, *SORBI_3004G149800*, is syntenic to both maize genes. Of the four maize genes, *mdr1* and *dng102* have the highest endosperm expression, and the expression of *mdr1* is more than double that of *dng102*, as indicated by expression of published mRNA-seq across multiple tissues (Stelpflug et al. 2016) (Figure 1D). Three of the four glycosylases, *mdr1*, *dng102*, and *dng103* had sufficient read coverage over maternal/paternal SNPs for analysis of imprinted endosperm expression in data from a recent study (Anderson et al. 2021). Of the three, *mdr1* has the most maternally biased expression, but not strong enough to be classified as a maternally expressed gene (MEG) (Figure 1E). One of the three, *dng102*, was paternally biased, but not strong enough to be classified as a paternally expressed gene (PEG).

We obtained two mutant alleles of *dng102*, the UniformMu insertion *mu1083641* (McCarty et al. 2005) in the C-terminal RRM-fold domain and a TILLING allele (Till et al. 2004) with a premature stop codon that replaces Q235, upstream of both the Endonuclease III domain and the RRM-fold domain. Neither allele produced a maternal R mottling phenotype, nor any obvious phenotype of any kind. However, based on PCR genotyping, we were unable to transmit either *dng102* mutant allele, neither maternally nor paternally, simultaneously with the *mdr1* mutation (Supplemental Figure S2). To screen larger numbers of kernels for possible rare events of double mutant transmission, we used GFP fluorescence linked to *Mdr1* and *Dng102* wild-type alleles in the Ds-GFP insertion collection (Li et al. 2013; Warman et al. 2020). By creating a *mdr1 dng102* double heterozygote with Ds-GFP insertions closely linked to each wild-type allele and crossing as male with non-GFP wild-type plants as females, we were able to screen thousands of kernels for paternal Ds-GFP inheritance (Figure 2A, B). All but six of 5221 resulting kernels were clearly fluorescent. PCR genotyping of four of the six ambiguous cases revealed that none had inherited both *dng102 mdr1* mutant alleles, and are likely explained by transgene silencing, pollen contamination, or recombination between Ds-GFP insertions and mutations. The remaining two kernels failed to germinate. We used both mutant alleles of *mdr1* for these crosses, the Kermicle allele (3786 resulting kernels) and EMS4-06835d (1435 resulting kernels). In control crosses with heterozygotes for both Ds-GFP insertions but homozygous wild-type for *Dng102* and *Mdr1*, 81 of 355 kernels lacked fluorescence. The 100% or near 100% inheritance of paternal Ds-GFP linked to wild-type *Mdr1* or *Dng102* raises the question of whether double mutant pollen leads to early seed abortion or fails to fertilize altogether. The organized structure of a maize ear allows an experimental means of testing this, as clusters of missing seeds would produce gaps in ears (Figure 2C). The production of evenly filled ears of fluorescent kernels from these crosses indicated that *mdr1 dng102* double mutant pollen (or earlier stage microgametophyte) have defects that prevent them from fertilizing (Figure 2B).

**Figure 2.**
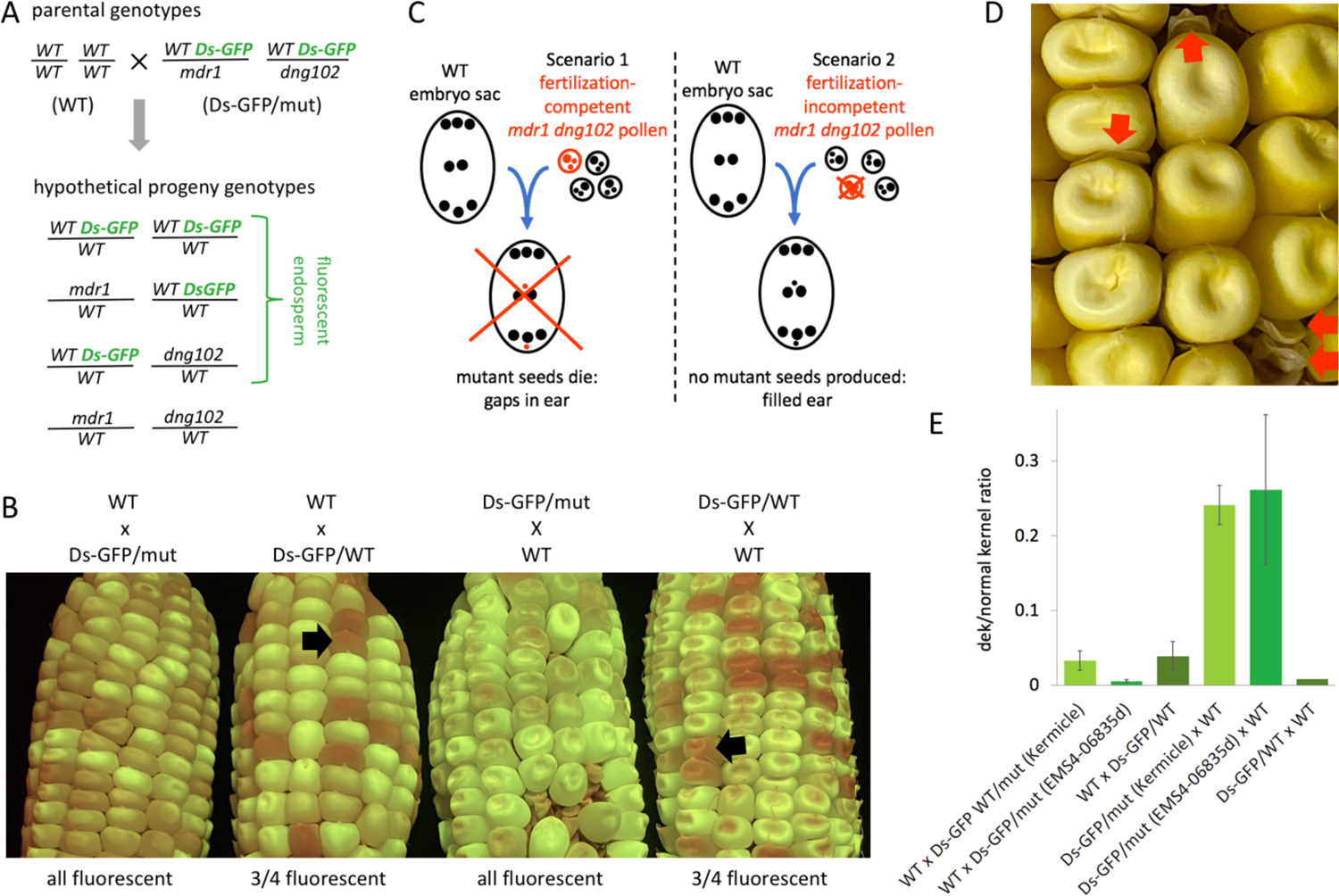
**A**, Schematic depiction of method to quantify transmission of *mdr1* and *dng102* mutant alleles using Ds-GFP insertions linked to the wild-type alleles. **B**, Example outcomes of crosses using Ds-GFP insertions. Ds-GFP/mut is the double heterozygous mutant of *mdr1* and *dng102* with the wild-type alleles linked to Ds-GFP insertions. Ds-GFP/WT is homozygous wild-type for both *Mdr1* and *Dng102*, but still heterozygous for the two Ds-GFP insertions. Black arrows indicate example non-fluorescent kernels. Fluorescence intensity depends in part on Ds-GFP dosage, from zero to four copies in triploid endosperm with insertions segregating at two loci. **C,** Schematic depiction of how well-filled ears indicates a pre-fertilization pollen defect. **D,** Defective kernel (dek) phenotype in seeds derived from a double heterozygous mutant mother plant crossed with wild-type pollen. **E,** Quantification of dek-like phenotype in the same crosses used to quantify Ds-GFP transmission. Error bars are standard errors of the means for each ear resulting from the crosses, except Ds-GFP/WT x WT, which was a single ear.

We also verified a requirement for maternal glycosylases by reversing the direction of the Ds-GFP crosses: non-GFP wild-type plants as females, *mdr1 dng102* double heterozygote with Ds-GFP insertions closely linked to each wild-type allele as males. All of 401 resulting kernels from the Kermicle *mdr1* allele were fluorescent, as were all of 202 kernels from the EMS4-06835d allele. In a control cross with heterozygotes for both Ds-GFP insertions but homozygous wild-type for *Dng102* and *Mdr1*, 71 of 240 kernels lacked fluorescence. In the maternal mutant but not wild-type crosses, the resulting ears had gaps, presumably corresponding to locations of double mutant megagametophytes (Figure 2D). Within these gaps, initiation of kernel development was often evidenced by visible pericarps of a range of sizes, similar to mutants with early collapsed defective kernel (dek) phenotypes (https://www.maizegdb.org/data_center/phenotype). The ratio of defective kernels to normal would be expected to be 1 defective to 3 normal if every double mutant megagametophyte produced a defective kernel. We found a ratio of closer to 1 to 4 (Figure 2E). These numbers should be interpreted cautiously, however, because in many cases it was difficult to distinguish very small pericarps from non-fertilization events, and they may have been hidden when wedged between normal kernels. In a control cross with a heterozygote for both Ds-GFP insertions but homozygous wild-type for *Dng102* and *Mdr1*, 71 of 240 resulting kernels lacked florescence and showed no evidence for a dek phenotype (Figure 2E). While the ear-to-ear frequency of defective kernels was variable, it was significantly lower in maternal mutant crosses compared to paternal mutant crosses for both the Kermicle and EMS4-06835d alleles (in both cases P-values < .00001, two-tailed Student t-tests). These results indicate that, unlike microgametophytes, megagametophytes that inherited both *mdr1* and *dng102* mutations are fertilization competent, but the resulting seeds die at an early stage in development.

### Identification of high-confidence sets of genomic regions that are demethylated in endosperm

To determine whether the *mdr1* mutant has detectable changes in DNA methylation relative to wild-type, we prepared and sequenced EM-seq libraries from homozygous wild-type and *mdr1* mutant developing endosperm 15 days after pollination (15-DAP), as well as the corresponding embryos. Based on the R mottling phenotype, we expected that the *r1* gene or regulatory sequence would be demethylated in endosperm with at least partial dependence on MDR1 activity. In the sequenced W22 genome and in other mottling-competent haplotypes, *r1* is not a single-copy gene, but a tandem duplicate gene complex. This complex includes at least one copy that is expressed in the aleurone layer of endosperm, part of whose 5’ UTR and entire promoter have been replaced by an approximately 400-bp CACTA transposon (Walker et al. 1995; May and Dellaporta 1998). The CACTA transposon, called the sigma region, is present in unmapped scaffold 282 of the W22 assembly immediately upstream of the gene *Zm00004b040676*, which encodes a MYC transcription factor homologous to *r1*. A second copy, *Zm00004b040677,* is annotated approximately 6 Kb upstream on the same scaffold, but with an N-gap region in place of a 5’UTR and promoter. *Zm00004b040676* itself contains three internal N-gap regions. We examined methylation in an 8-Kb region of scaffold 282. While N-gaps and non-unique sequence prohibit EM-seq read mapping across most of the region, reads do map to an approximately 1-Kb region including the 5’ end of *Zm00004b040676* and sigma region and reveal increased methylation in *mdr1* mutant relative to wild-type endosperm. Unlike in endosperm, the sigma region was highly methylated in other wild-type plant tissues including embryo (Figure 3A and Supplemental Figure 3).

**Figure 3.**
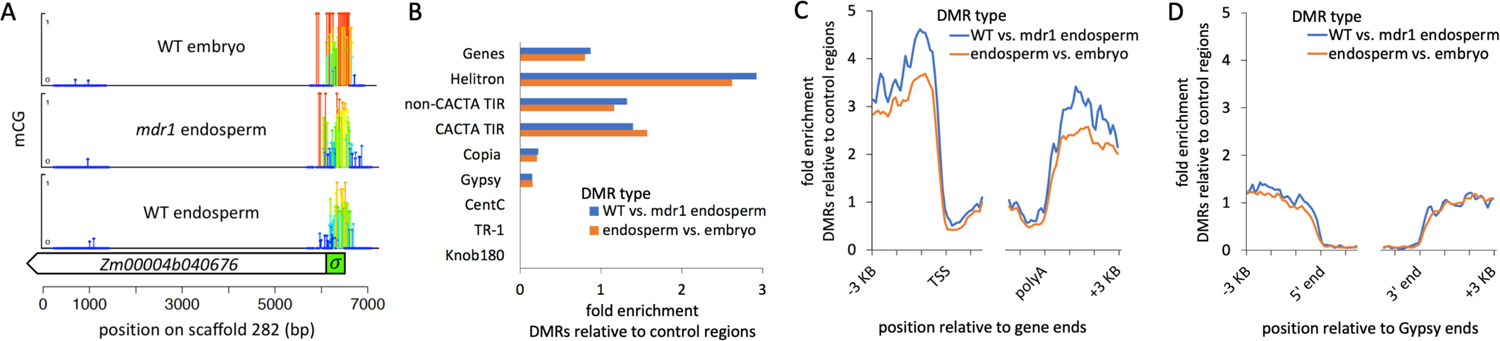
**A**, Differential methylation of an *r1* gene in embryo, endosperm, and *mdr1* mutant endosperm. mCG values are shown at single-base level resolution over a 8-Kb region including the majority of an *r1* homolog in W22 (*Zm00004b040676*). Both height and color of lollipops indicate mCG values (darkest blue = zero mCG). Regions without lollipops indicate lack of read coverage, either because of multi-copy sequence or large N gaps. **B**, DMR enrichment in different genetic elements. Enrichment is the proportion of DMRs that overlap each element by at least 50% of their length (100 bp) divided by the proportion of control regions that do. Control regions are the 200-bp regions that were eligible for identifying DMRs based on read coverage and number of informative cytosines. **C,D,** Enrichment for DMRs on 100-bp intervals relative to gene ends (C) or Gypsy ends (D). Gypsies with solo LTRs were excluded from this analysis. For genes, TSS = Transcription start site, polyA = polyadenylation site.

We also used these methylomes to search genome-wide for differentially methylated regions (DMRs) between homozygous wild-type and *mdr1* mutant 15-DAP endosperm (“WT vs. mdr1 endosperm”). In parallel, we also searched for DMRs between wild-type endosperm and wild-type embryo (“endosperm vs. embryo”. In order to identify regions with high-confidence differential methylation, we used a conservative approach that required differential methylation of both mCG and mCHG over non-overlapping 200-bp regions. First we identified the set of all regions that were eligible for differential methylation analysis based on their having at least 3X average coverage of at least five CGs and five CHGs in both tissues. From this set of eligible regions, we identified hyperDMR as having a greater than twofold relative increase in percent methylation (value2/value1 > 2) and a greater than 20% relative increase in methylation (value2 – value1 > 0.20) for both mCG and mCHG independently. HypoDMRs satisfied the opposite requirements. We expected these strict criteria for defining DMRs would not produce exhaustive sets but would produce high-quality representative set of DMRs for each of the comparisons. We used the entire sets of eligible regions as controls regions for DMRs, since they were subject to the same biases in terms of ability to map EM-seq reads and containing informative CG and CHGs. Such controls are essential in a large repetitive genome like maize, where even with high coverage, much of the genome is inaccessible to methylation measurements by short reads.

In wild-type vs. mdr1 mutant endosperm, we found 18,464 hypoDMRs and 563 hyperDMRs. In wild-type endosperm vs. wild-type embryo, we found 52,919 hypoDMRs and 373 hyperDMRs. For comparison, we also prepared and analyzed methylomes from wild-type and mutant premeiotic tassel and wild-type and mutant mature endosperm. The *mdr1* mutant produced large numbers of hypoDMRs in mature and 15-DAP endosperm, but not in tassel or embryo (Supplemental Figure S4). Given that we expected only the hypoDMRs to be a direct result of DNA glycosylase activity in endosperm, we excluded hyperDMRs from further analysis; and for brevity we refer to the hypoDMRs as simply DMRs.

### Regions that are demethylated in endosperm are depleted of retrotransposon and tandem repeats

We examined the genomic locations of the DMRs produced in both the WT vs. mdr1 endosperm comparison and endosperm vs. embryo comparison. Both sets behaved similarly in being generally depleted from the most abundant repetitive elements in the genome (retrotransposon and tandem repeats) but were mildly enriched flanking genes (Figure 3B-D and Supplemental Table S2). The large number of elements in each group means that every enrichment, even for TIR DNA transposons with fold enrichment values a little above 1, is statistically significant. Only Helitrons, however, were enriched more than twofold. The complete absence of DMRs from the tandem repeats *Knob180*, *CentC* and *TR-1* was not due to an inability to map reads because we identified thousands of control regions in them (Supplemental Table S2). The enrichment of DMRs upstream and downstream of genes raises the possibility that they might be specifically demethylated at heterochromatin boundaries to extend the length of euchromatin domains in endosperm. To test this, we examined the distribution of distances between DMRs and the closest euchromatin regions, using regions low in mCHG in embryo as the defining feature of euchromatin. In most maize tissues at most loci, euchromatin is stably depleted of mCHG across development (Oka et al. 2017; Crisp et al. 2020; Hufford et al. 2021). Consistent with both heterochromatic and euchromatic activity of DEMETER in Arabidopsis endosperm (Frost et al. 2018), almost half of the DMRs were greater than 2 Kb from the nearest euchromatin, indicating that MDR1 also effectively demethylates deep in heterochromatin, not just at its edges (Supplemental Figure S5A).

Motif analysis did not reveal anything significant except ones mainly consisting of strings of A’s and T’s. Consistent with that, DMRs had lower GC content than control regions: 41.5% in DMRs and 49.0% in control regions for WT vs. mdr1 endosperm DMRs (Supplemental Figure S5B). An analysis of transposons at family-level resolution confirmed the depletion (or nearly neutral status) of DMRs for nearly all families, with one notable exception: Over a thousand copies of Helitrons from the DHH00002 family overlapped DMRs (1060 copies in WT vs. mdr1 endosperm and 1748 in endosperm vs. embryo), with about 8-fold higher than expected frequency based on control regions. 16 other TE families also were enriched for DMRs relative to control regions, but these were mainly low copy families with only a handful of copies (Supplemental Figure S5C). Including the abundant DHH00002 family, these 17 families overlapped with 11.7% of the WT vs. mdr1 endosperm DMRs (a 7.3 fold enrichment for DMRs relative to control regions).

The scarcity of DMRs in gene bodies, especially near their transcription start sites and polyadenylation sites (Figure 3C), is not surprising given the normal absence of mCG and mCHG from these regions, which by definition makes demethylation impossible. An alternative way to think about DMR enrichment is relative to the subset of regions that have the potential to be methylated. To explore this, we defined methylated control regions as ones with both mCG and mCHG values of at least 0.2 in at least one of the two methylomes in each comparison. When we compared DMRs with this set of methylated control regions rather than all control regions, we measured an approximately 8-fold enrichment for DMRs internal to both ends of genes (Supplemental Figure S6).

### The *mdr1* mutant reduces both methylation and siRNAs in endosperm

Measuring methylation levels in and flanking DMRs confirmed their enrichment near euchromatin and revealed that kilobase-scale regions extending beyond the 200-bp regions defined by our method were detectably demethylated (Figure 4A-F and Supplemental Figure S7). In the *mdr1* mutant endosperm, mCG and mCHG were partially restored to non-endosperm states in DMRs. At a genome-wide level, however, *mdr1* had little effect, as evidenced by the similar levels of methylation in control regions in mutant and wild-type (Figure 4A-C). mCHH, which indicates RdDM activity in maize endosperm as well as other tissues (Fu et al. 2018), was low in DMRs independently of *mdr1* or tissue. A prediction of the siRNA transfer model is that increase methylation in *mdr1* mutant in endosperm would be accompanied by decrease in methylation in embryo. This was not the case: methylation was either the same or slightly increased in *mdr1* mutant embryo relative to wild-type.

**Figure 4.**
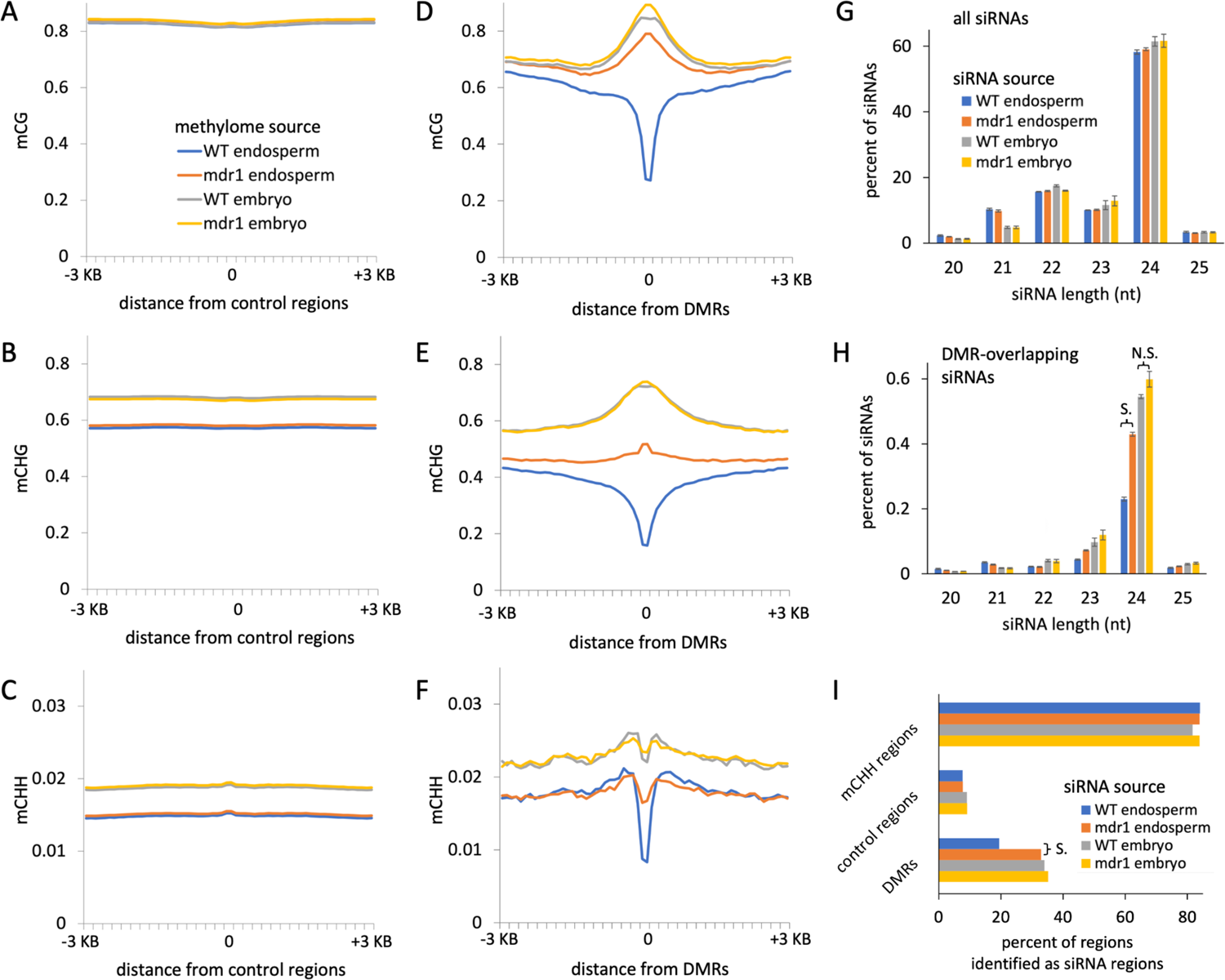
**A-C**, mCG, mCHG, and mCHH profiles centered on control regions associated with wild-type vs. mdr1 endosperm DMRs. Methylation values are averages of 100-bp intervals within and up to 3 Kb on either side of each 200-bp region. **D-F,** Same as A-C, except methylation centered on wild-type vs. mdr1 DMRs instead of control regions. **G**, Length distributions of all siRNAs. Error bars are standard errors of the means for biological replicates. **H**, As in G, except only siRNAs that overlapped at least 90% of their length with wild-type vs. mdr1 endosperm DMRs. “S.” indicates significant difference: P-value < .00001, two-tailed Student t-test. “N.S.” indicated not significant: P-value = .06. **I**, Regions categorized by whether they produce siRNAs. An “siRNA region” is any that had at least 50 bp spanned by siRNAs of the indicated source tissues. All libraries were subsampled to 20 million siRNA reads. DMRs and control regions are from the wild-type vs. mdr1 endosperm comparison. mCHH regions are 200 bp regions with an average mCHH value of at least 0.2 in wild-type endosperm. “S.” indicates significant difference (P-value < .0001 two-tailed chi-square with Yates correction).

In Arabidopsis, demethylation of heterochromatin in pollen vegetative cells corresponds to expression of 21 and 22nt siRNAs from LTR retrotransposons (Creasey et al. 2014; McCue et al. 2015; Borg et al. 2021). A similar phenomenon occurs in maize embryos, in response to partial demethylation of heterochromatin in mutants of DDM1-type nucleosome remodelers or chromomethyltransferases (Fu et al. 2018). Even though MDR1 activity appears to inhibit the little mCHH that is present in endopsperm DMRs (Figure 4A-F and Supplemental Figure S7), it is possible that siRNAs produced by endosperm demethylation could direct other chromatin modifications or function post-transcriptionally (Parent et al. 2021). To directly test the effect of the *mdr1* mutant on siRNA expression, we sequenced small RNA libraries in the same tissues that we used for methylome analyses. We classified all small RNAs of 20 to 25 nt in length that did not overlap at least 90% of their lengths with tRNAs, ribosomal RNAs, or miRNAs as siRNAs. The length distribution of total siRNAs did not change between wild-type and mutant endosperm, but there was a nearly twofold increase in the abundance of 24-nt siRNAs that overlapped with DMRs in *mdr1* mutant (Figure 4G,H). As with methylation, *mdr1* partially reverted endosperm to an embryo or tassel-like 24-nt siRNA profile at DMRs (Supplemental Figure 8). This pattern was evident whether including just uniquely mappable siRNAs, or also including multi-mapping ones. The *mdr1* mutant increased not only the total abundance of siRNAs, but also the number of DMRs that produced detectable levels of siRNAs (Figure 4I). The number of DMRs that produced detectable siRNAs increased from 19% of DMRs in WT to 33% in *mdr1* mutant (p-value < .0001 two-tailed chi-square with Yates correction). These data, combined with methylation data, indicate that MDR1 activity in endosperm inhibits rather than promotes siRNA expression in endosperm.

### Endosperm DMRs imprint expression of a subset of DMR-containing genes and transposons

Since endosperm inherits two copies of each maternal genome and one of each paternal genome, the expression of maternal transcripts would be expected to account for two thirds of the total transcript in endosperm. Reciprocal hybrid crosses, where maternal and paternal alleles can be differentiated by SNPs, provide a means to test this. The maternal ratio of 0.67 in maize endosperm indeed corresponds to twice as high expression of alleles that are inherited maternally than paternally (Waters et al. 2011; Zhang et al. 2011; Anderson et al. 2021). The majority of genes that overlapped with DMRs and were expressed in endosperm showed little or no evidence for imprinting (Figure 5A). While DMRs did not strongly predict imprinting, imprinting did strongly predict DMRs. For both comparisons, there was an approximately 9-fold increase for overlap with DMRs among MEGs relative no non-imprinted genes (Figure 5B). Both comparisons were statistically significant (P-value < .00001, two tailed chi-square test). We also tested whether DMRs in the 2 Kb flanking genes correlated with imprinting, but did not find any clear pattern (Figure 5B). Endosperm expression in general did not predict overlap with DMRs (Figure 5C). DMRs in MEGs were preferentially located towards their 5’ ends (Figure 5D) while DMRs in genes as a whole had a more balanced representation at both ends (Figure 5E).

**Figure 5.**
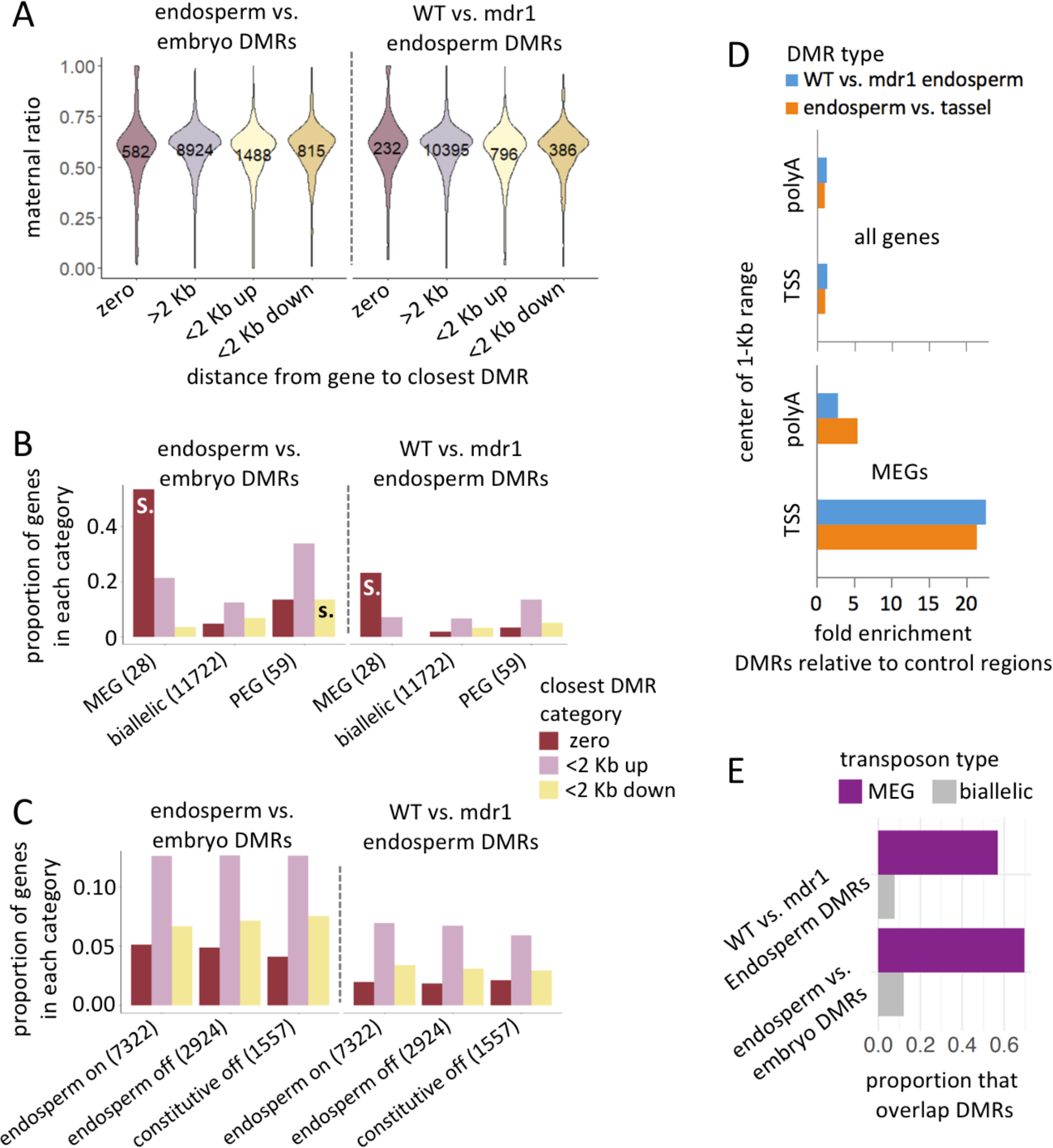
**A**, Genes that overlap endosperm DMRs are skewed toward maternal expression but are still predominantly biallelically expressed. A value of 1 means completely maternal expression and 0 means completely paternal (Anderson et al. 2021). Genes are categorized according to distance from endosperm DMRs. Numbers of genes in each category are indicated. **B**, MEGs preferentially overlap endosperm DMRs. Genes are categorized according to distance to closest DMR (by color) and according to imprinting status (X-axis labels, including total number of genes in each imprinting category). MEGs are defined by a maternal ratio of greater than 0.9, PEGs as less than 0.1, biallelic as intermediate values. Only genes with SNPs and sufficient coverage to make imprinting calls are included. Significant differences between imprinted and biallelic sets are indicated by “S.” (P-value < .00001, two tailed chi-square test), or “s.” (P-value < .01, two tailed chi-square test). The larger proportions for endosperm vs. embryo are because of the larger number of DMRs (52191, compared to 18464 in WT vs. mdr1 endosperm. **C**, Endosperm gene expression is not correlated with DMRs. Genes are categorized according to distance to closest DMR (by color) and according to expression status (X-axis labels, including total number of genes in each expression category). “Endosperm on” means expressed in endosperm, “endosperm off” means not expressed in endosperm but expressed in other tissues, and “constitutive off” means not expressed in endosperm and not expressed in other tissues. Data from Stelpflug et al. 2016. No differences between endosperm on and off categories were significant (P-value > .01, two tailed chi-square test. **D**, MEGs are enriched for endosperm DMRs near their 5’ ends. DMR enrichment is the number of DMRs that overlap with a 1-Kb range either centered on transcription start sites (TSS) or polyadenylation sites (polyA) of MEGs normalized by the number of control regions that overlap the same range. **E**, Maternal imprinted transposons have a high frequency of overlap with endosperm DMRs. MEG-type transposons (95 total) have an maternal ratio value greater than 0.9, PEG as less than 0.1, and non-imprinted (1773 total) as all intermediate values. The Y-axis indicates the proportion of transposons that overlap endosperm DMRs from each of the two comparisons. In both, MEG-type transposons are significantly enriched for DMRs over non-imprinted ones (p-value < 0.01, binomial test). Only one PEG-type transposon overlapped with DMRs and is not shown.

The majority of transposon copies with sufficient expression in endosperm and with variants to allow distinguishing parent of origin also have maternal ratio values near 0.67 (Anderson et al. 2021). However, we found that the 5% of expressed transposons with MEG-type expression are 4.8-fold more likely to overlap endosperm DMRs than the endosperm-expressed but non-imprinted set (Figure 5E). Only three transposon copies (0.16% of the endosperm-expressed set) had PEG-type expression, too few for a meaningful analysis. These results indicate that demethylation in endosperm-expressed genes usually has little or no effect on genomic imprinting, but in specific situations demethylation can cause maternal-specific expression of both genes and transposons.

## DISCUSSION

In Arabidopsis, DEMETER is required for normal development not just in endosperm, but in the pollen vegetative cell and non-reproductive cell types as well (Schoft et al. 2011; Borg et al. 2021; Khouider et al. 2021; Kim et al. 2021). In the case of pollen vegetative cell, developmental phenotypes are determined by gene regulation rather than transposon activation. Here we have demonstrated in maize that it is essential to have a functional copy of at least one of the DNA glycosylase genes *mdr1* or *dng102* prior to fertilization in both the maternal and paternal gametophytes, based on inability to transmit two mutant alleles simultaneously (except to produce seeds that die early in development in the case of maternal transmission) (Figure 2).

Imprinted gene expression in maize is clearly linked to demethylation of maternal alleles (Zhang et al. 2014). Such developmentally dynamic methylation in endosperm gene regulation is an exception to the general rule of mCG and mCHH methylation being stable (Oka et al. 2017; Crisp et al. 2020). In multiple species, demethylation of transposons near MEGs is clearly connected to their maternal expression (Gehring et al. 2009; Hsieh et al. 2011; Hatorangan et al. 2016; Pignatta et al. 2018; Rodrigues et al. 2021). A prediction from these observations is that a genome like maize, with more frequent transposons near genes, would have larger numbers of imprinted genes. This, however, is not the case. In addition, we found that endosperm DMRs were only weakly associated with transposons, with the exception of many Helitrons (Figure 3). Transposon families that were enriched for endosperm DMRs accounted for about a tenth of total endosperm DMRs. The vast majority of the enriched transposon copies were of the DHH00002 Helitron family. It is clear both from our findings and prior work on other species that DNA glycosylases demethylate repetitive elements and that their demethylation can cause imprinted gene expression in endosperm [reviewed in (Anderson and Springer 2018; Batista and Köhler 2020)]. Our results are consistent with this, but further indicate that demethylation of transposons is not the major function of DNA glycosylases in maize endosperm.

It has been proposed that transposon demethylation would make sense in the context of demethylation being part of a transposon silencing mechanism, where demethylation promotes siRNA expression that transfer into egg or embryo (Ibarra et al. 2012; Bouyer et al. 2017). Both our siRNA expression and DNA methylation data argue against this. The *mdr1* mutant had no decrease in methylation nor siRNAs in embryo, but it did have a clear increase in siRNAs in endosperm (Figure 4). Recent work in *Brassica rapa* also found no evidence to support the idea of endosperm siRNAs being transferred to embryo (Chakraborty et al. 2021). Related to this, the preferential demethylation near genes in endosperm suggests the possibility of interaction with RdDM, which is strongly enriched near genes (Gent et al. 2014; Fu et al. 2018). However, the low mCHH and siRNA abundance in demethylated regions in endosperm suggest otherwise.

Consistent with what is known about glycosylase activity in other plants, MEGs were strongly enriched for overlapping endosperm DMRs, especially in the 5’ end of genes. However, the majority of genes that were expressed in endosperm and contained endosperm DMRs were not imprinted (Figure 5). This raises two possibilities: First, that the effect of MDR1 and (likely DNG102 as well) is not limited to one parent. The idea that glycosylases act post-fertilization on both parental genomes is consistent with the biparental expression of both *mdr1* and *dng102* in endosperm long after the two parental genomes have merged (Figure 1). If glycosylases act on both genomes in endosperm, both maternal and paternal alleles would have reduced methylation relative to non-endosperm control tissue. Testing this will require comparison of hybrid methylomes of endosperm and non-endosperm tissues, where SNPs allow for distinguishing of parental alleles. The second possibility is that endosperm demethylation has an additional function independent of transcriptional regulation. Endosperm may be unusual in its rapid rate of cell division, which partly takes place in a syncytium [reviewed in (Becraft and Gutierrez-Marcos 2012)]. At least in Arabidopsis, endosperm also has unusual distributions of its chromosomes and less condensed chromatin then typical nuclei (Baroux et al. 2007; Baroux et al. 2017; Yadav et al. 2021). Removal of DNA methylation from select regions of the genome may facilitate rapid DNA replication or related chromatin changes in endosperm. While *mdr1* null mutants effectively behave as partial-loss-of-function due to complementation by its homolog *dng102*, the viability of *mdr1* endosperm and pollen provides an experimental resource for investigating these and related phenomenon.

## METHODS

### Candidate *mdr1* mutation mapping

As shown in Supplemental Figure S1A, homozygous *mdr1* plants derived from stock X336J (from the Maize Genetics Cooperation Stock Center, http://maizecoop.cropsci.uiuc.edu/) were crossed with the W22 sequenced stock (Springer et al. 2018) to create *mdr1/Mdr1* heterozygotes. These were crossed as females with *mdr1* homozygous males derived from stock X336J to produce progeny segregating homozygous and heterozygous *mdr1*. All plants were homozygous for the *R1-r:standard* haplotype which is capable of mottling. Approximately 500 putative *mdr1* homozygotes were selected based on kernel mottling and planted in five batches of 100 each. Single leaf tips from each seedling were combined for each of the five batches, and a single DNA extraction performed on each batch, followed by Illumina library preparation with a KAPA HyperPrep Kit (KK8502). Three batches were sequenced single-end and two sequenced paired-end, all with 150-nt reads. A single batch of five W22 leaf tips was also processed in the same manner as a homozygous *Mdr1* control and sequenced single-end. Adapter sequences were trimmed from all reads using cutadapt 1.9.1 (Martin 2011) with adapter sequence AGATCGGAAGAGC for both forward and reverse reads and parameters -q 20 -O 1 -m 100. Trimmed reads were aligned to the W22 reference genome (Springer et al. 2018) using BWA mem version 0.7.15 (Li and Durbin 2009) with the -M parameter.

To generate a set of high-confidence variants, duplicates were marked using Picard MarkDuplicates (v2.16.0 http://broadinstitute.github.io/picard). Downstream variant calling was performed using software in the GATK package (v3.8.1) and the GATK best practices (Van der Auwera et al. 2013). In particular, indels were realigned using RealignerTargetCreator and IndelRealigner, and an initial round of SNP calling was performed using HaplotypeCaller and GenotypeGVCFs. The resulting vcf files for the W22 bulk and the combined *mdr1* bulks were filtered to remove variants that were present in the W22 bulk using the BEDTools subtract tool version 2.26.0 (Quinlan and Hall 2010) and stringently filtered to remove low quality score variants using VCFtools 0.1.15 (Danecek et al. 2011) with the -minQ 200 parameter. The resulting vcf was used as a training set for base recalibration of the W22 bulk and *mdr1* combined bulks using BaseRecalibrator, and HaplotypeCaller/GenotypeGVCFs was used to call variants in W22 and *mdr1*. VariantsToTable was used to convert vcf data into a readable table for analysis in RStudio.

The mapping scheme predicted that variants across the genome would have a frequency of 0.75 non-W22 variants (alternative alleles). Since we were looking specifically for a region was near homozygous for alternative alleles in the *mdr1* bulks, the following additional criteria were imposed to identify putative variants linked to *mdr1* in the vcf table: 1) An alternative allele frequency in the W22 bulk of zero, with a read depth of at least two but no more than 15. 2) A read depth in the *mdr1* combined bulks of at least five but no more than 50. 3) A normalized Phred-scaled likelihood for homozygous reference alleles of at least 400 and for homozygous alternative alleles of zero in the *mdr1* combined bulks. Filtering the vcf table in this way selected for near-homozygous alternative alleles across the genome in the *mdr1* bulks, with an average of 96% alternative allele frequency. A single ∼20 MB region (chr4:216965546-237367197) stood out, with a ∼98.5% alternative allele frequency. A second method of analyzing the GATK variants was also used to identify the *mdr1* candidate region. The vcf table was filtered to exclude all variants but those with a GenotypeQuality of at least 20 in both genotypes, depth of coverage between 5-14 in the W22 bulk, a depth between 6-49 in the combined *mdr1* bulks, and variant frequencies above 0.75 in the *mdr1* bulks. Based on the mapping scheme, this is expected to remove the majority of technical artifacts. As with the first method, this singled out the same region on chromosome 4 as the most homozygous in the genome, with variant frequencies near 1, as shown in Supplemental Figure 1B. A comparison of variant density by genomic position revealed that a large region of chromosome 5 as well as a few small regions elsewhere in the genome of the *mdr1* mutant stock X336J were not shared with the sequenced W22 genome (Supplemental Figure S1C). The tip of chromosome 4, however, had few variants and was clearly shared between both genomes.

### Mutant genotyping

An 867-bp region of *mdr1* was amplified by PCR with primers W22-Ros1-F1 and W22-Ros1-R1 and digested with TaqI-v2 (NEB #R0149S). The different *mdr1* alleles could be distinguished by the pattern of restriction fragments due to the presence of SNPs that created or removed TaqI restriction sites in different genetic backgrounds (Kermicle’s *mdr1* allele or its progenitor in W22, the *EMS4-06835d* allele or its progenitor in B73, and the allele linked to the Ds-GFP insertion *R128F11* in A188). A single TaqI restriction site in B73 yields two fragments of 382 and 485 bp. A second site in A188 yields three fragments, 107, 275, and 485 bp. Absence of any sites in W22 yields an intact amplicon of 867 bp. In cases where the wild-type and mutant were in the same genetic background or when there was a possibility of recombination between *R128F11* and *mdr1*, the *mdr1* mutation was genotyped by Sanger sequencing of the amplicon.

Kermicle’s *mdr1* allele reduced the string of seven A’s to six in the W22 amplicon subsequence AGTTTACGAAAAAAAGTGCTTTC. Primers are as follows:

>W22-Ros1-F1 TTAGAACAAGGCGACTTTTCACC

>W22-Ros1-R1 AGGAACTCGGTAGCTCTTCT

To genotype the *dng102-Q235 allele*, A 569-bp region of *dng102* was amplified by PCR with JIG-339 and JIG-340 primers and digested with Hpy188I (NEB # R0617S). This amplicon has two Hpy188I sites in wild-type, but the mutation that converts Q to a stop codon at amino acid position 235 removes one of them such that digestion of the wild-type amplicon yields three fragments of 317, 156, and 96 bp, but the *dng102-Q235* allele yields two fragments of 473 and 96 bp. Primers are as follows:

>JIG-339 TCCAAAGCCTCCCAAAGAAA

>JIG-340 GCATATGATCGACCGAGCTG

To genotype the *dng102 mu1083641* allele, short regions were amplified using EoMumix3 primers along with either of two *dng102* primers, JIG-337 or JIG-338, which were also used together to amplify the wild-type locus. Primers are as follows:

>EoMumix3 GCCTCYATTTCGTCGAATCCS

>JIG-337 CTGGCAGGAAAATGCAACCT

>JIG-338 GCAATTGCGCAAGCTTAATG

To genotype the Ds-GFP insertion *R153F01* linked to *Dng102*, two primer combinations were used: the Ds-GFP primer JSR01dsR with the flanking primer JIG-368, and the Ds-GFP primer JGp3dsL with the flanking primer JIG-369. JIG-368 and JIG-369 were also used as a pair to amplify the wild-type locus. Primers are as follows:

>JSR01dsR GGGTTCGAAATCGATCGGGATA

>JGp3dsL

ACCCGACCGGATCGTATCGG

>JIG-368 CGGAGTCATTCTTGCTCTTCT

>JIG-369 AGGACGAGACCTTAGTCCTTAT

### Relationships between DNA glycosylases

MDR1-like DNA glycosylases in *Arabidopsis*, rice, sorghum, and maize were identified by aligning the amino acid sequence of MDR1 against proteins of each species using Gramene BLAST (Tello-Ruiz et al. 2021). TAIR10, ASM465v1, Sorghum_bicolor_NCBIv3, and Zm-B73-REFERENCE-NAM-5.0 were used as reference sequences. All homologs that were annotated to have both the Endonuclease III and RRM-fold domains were included for comparison, with the exception of ROS1B in rice, which was included despite lacking an RRM-fold domain. Transcript IDs for all genes as well as percent amino acid identities are listed in Supplemental Table S1. Amino acid identities were calculated using the Geneious multiple alignment tool (version 10.1.2; http://www.geneious.com) using global alignment type, identity cost matrix, gap open penalty of 12, and gap extension penalty of 3. The percent amino acid identities in Figure 1B and Supplemental Table S1 are the number of identical amino acids divided by the sum of the number of identical amino acids and number of different amino acids over the entire protein length. The protein tree in Figure 1B was produced using the Genieous tree builder tool using global alignment type, identity cost matrix, Jukes-Cantor genetic distance model, UPGMA tree build method, gap open penalty of 12, and gap extension penalty of 3.

Synteny between sorghum and maize genes was determined by presence of collinear genes as revealed by the CoGe GEvo tool (https://genomevolution.org/coge/GEvo.pl). For these analyses, Sorghum glycosylase genes were used as references and maize glycosylase genes as queries. 4-Mb regions of genomic sequence centered on each maize gene was aligned to 4-Mb regions of sorghum genomic sequence centered on each sorghum gene. Sorghum non-CDS sequence was masked, maize unmasked.

### Ds-GFP screening for mutation transmission

Stocks containing Ds-GFP alleles (R128F11 at chr4:229537027 and R153F01 at chr5:165072386) were obtained from the Maize Genetics Cooperation Stock Center. Double heterozygous plants (*R128F11 Mdr1/mdr1*; *R153F01 Dng102/dng102*) were reciprocally crossed with wild-type Oh43 inbred plants without Ds-GFP insertions, as shown in Figure 2A. For *mdr1* mutant, both the Kermicle and EMS4-06835d alleles were used; for *dng102*, just *dng102-Q235*. GFP fluorescence in kernels was revealed under visible blue light using a Dark Reader Hand Lamp and Dark Reader Glasses (Clare Chemical Research #HL34T) similar to previous methods, but on a smaller scale without automated kernel counting (Warman et al. 2020). The *dng102-Q235* allele in 04IA-B73PS-055_C4 in B73 was backcrossed once into non-mutagenized B73, selfed twice, then crossed three times into either W22 or W22-related stocks before crossing with Ds-GFP stocks. EMS4-06835d in B73 was crossed twice into W22 before crossing with DsGFP stocks.

### Harvesting plant materials

Four organs were harvested for EM-seq and small RNA-seq library preparations: 1) Developing endosperm 15 days after pollination (15-DAP) were collected by removing embryos, pericarp, and nucellus with forceps. 10 endosperms were combined for each biological replicate, three replicates of homozygous *mdr1* (Kermicle allele in W22), and of homozygous wild-type (sequenced W22 inbred stock). Each replicate was derived from endosperms from a single ear. The homozygous mdr1 plants were derived by introgressing the W22-related *mdr1* homozygous stock derived from X336J into the sequenced W22 inbred stock six times followed by crossing two *mdr1* heterozygous siblings together. 2) 15-DAP embryos paired with the above endosperms were also collected. They were approximately 3.5 mm in width and 5 mm in length at this stage. 15 embryos were combined for each biological replicate, from the same ears as the endosperm. 3) Mature endosperm was separated from pericarp with forceps after soaking in water for 20 minutes and separated from embryo by dissection with a razor blade. Three replicates each of *mdr1* homozygous and *Mdr1* homozygous siblings were included, where each replicate was derived from a single kernel. The heterozygous parents that were crossed to produce these plants were the product of a cross between the sequenced W22 inbred stock and the W22-related *mdr1* homozygous stock derived from X336J. 4) Premeiotic tassels were collected from plants at approximately the 15-leaf stage, when tassels were between 0.8 and 2.3 cm in length. Three replicates consisting of two or three individual tassels from *mdr1* homozygous and wild-type homozygous siblings were included. These plants were grown from siblings of the ones used for mature endosperm collections.

### Preparation of Enzymatic Methyl-seq (EM-seq) sequencing libraries

DNA was extracted using IBI plant DNA extraction kits (#IB47231) after grinding tissue to a fine powder (in liquid nitrogen for tassel, 15-DAP endosperm, and 15-DAP embryo), and NEBNext® Enzymatic Methyl-seq Kits (#E7120S) used to prepare libraries. The input for each library consisted of 200 ng of genomic DNA that had been combined with 1 pg of control pUC19 DNA and 20 pg of control lambda DNA and sonicated to fragments averaging ∼700 bp in length using a Diagenode Bioruptor. The protocol for large insert libraries was followed. Libraries were amplified with 4 or 5 PCR cycles. Libraries were Illumina sequenced using paired-end 150 nt reads. Read counts, and accession numbers, and conversion measures are listed in Supplemental Table S3.

### Preparation of small RNA sequencing libraries

The same materials used for preparation of EM-seq libraries were also used for preparation of small RNA sequencing libraries, except mature endosperm. RNA was extracted using mirVana miRNA isolation kits (Thermo Fisher Scientific #AM1560) using the total RNA method for endosperm, embryo, and leaf; and small RNA enrichment method for tassel. Plant RNA Isolation Aid (Thermo Fisher Scientific #AM9690) was added at the lysis step except for embryo samples. Small RNA sequencing libraries were prepared using the NEXTflex Small RNA-Seq Kit v3 (PerkinElmer #5132-05) using the gel-free size selection & cleanup method. ∼500 ng small RNA enriched RNA was used as input for each tassel library and 550-2000 ng total RNA used for other organs. Libraries were amplified with 13 to 15 cycles of PCR. Libraries were Illumina sequenced using paired-end 150 nt reads. Mapped read counts and accession numbers are listed in Supplemental Table S4.

### Transposon, tandem repeat, and gene annotations

Transposon annotations (W22.allTE.withSolo_10Jan18.gff3.gz) were downloaded from https://ftp.maizegdb.org/MaizeGDB/FTP/Zm-W22-REFERENCE-NRGENE-2.0/. The transposon annotations are based on a conservative identification method using intact known structural features (Springer et al. 2018; Anderson et al. 2019a). Retrotransposons having solo LTRs were distinguished from those having both LTRs in these annotations by “Zm00004bS” in their names. Superfamilies were identified by their three-letter codes, e.g., RLG. Locations of tandem repeats in the genome were determined by mapping dimerized consensus sequences for *CentC*, *knob180*, and *TR-1* (Gent et al. 2017; Liu et al. 2020) to the genome using Bowtie2 (Langmead and Salzberg 2012) (version 2.4.1) with the --local parameter. All tandem repeat loci that were within 50 bp of each other were merged with the BEDTools merge tool with the -d 50 parameter. Following this, all loci less than 100 bp in length were removed. Locations of tRNA were identified by mapping tRNA sequences from the Genomic tRNA Database (Chan and Lowe 2016) (http://gtrnadb.ucsc.edu/GtRNAdb2/genomes/eukaryota/Zmays7/zeaMay7-tRNAs.tar.gz) to the genome using Bowtie2 with the --local parameter. All loci less than 70 bp in length were removed. Locations of 5S ribosomal RNA were identified by mapping NCBI Reference Sequence XR_004856865.1 to the genome using Bowtie2 with the --local parameter. All loci less than 100 bp in length were removed. Locations of nucleosome organizing region (NOR) ribosomal RNAs were identified first in the B73 v5 reference genome (Hufford et al. 2021) by blasting the NCBI reference 28S ribosomal RNA sequence XR_004853642.1 as a query with default parameters for a high similarity blast except a word size of 15 and an e-value of 1e-295 on maizeGDB. This identified an approximately 12.4-Kb tandemly repeated NOR sequence. One complete repeat sequence (chr6:16838268-16850655) was then blasted to the W22 genome using BLAST+ with default parameters except -evalue 1e-100. All tandem repeat loci that were within 50 bp of each other were merged with the BEDTools merge tool with the -d 50 parameter. Locations of miRNAs were identified by mapping Zea mays mature and hairpin miRNA sequences downloaded from The miRBase Sequence Database -- Release 22.1 (Kozomara et al. 2019) to the W22 genome. First, perfectly matching loci were identified by mapping mature miRNA sequences to the genome using Bowtie2 with the -a and --very-sensitive parameters followed by grep -v ‘NM:i:1’. Mature miRNA sequences that did not have a perfect match to genome were then identified by mapping with the --very-sensitive parameter. Locations of miRNA larger hairpins were identified by mapping miRNA hairpin sequences to the genome with the -a and --local parameters after merging overlapping alignments using the BEDTools merge tool, all hairpin loci less than 60 bp in length were removed. Only mature miRNA alignments that overlapped their full length with hairpin loci were retained as miRNA loci. The location of the sigma region associated with *r1* in the W22 genome was identified by MaizeGDB BLAST using sequence from GenBank accession AF106323.1 as query (Walker et al. 1995). The scaffold carrying the sigma region and two *r1* genes, scaffold 282, is also named LWRW02000294_1.

W22 v2 gene annotations (Springer et al. 2018) were compared to B73 v4 gene annotations (Jiao et al. 2017) to identify genes that were annotated in both genomes. Only this high-confidence set of W22 gene annotations was used in all analyses. MEG and PEG status of genes in W22 were determined as previously described using reciprocal crosses between W22 and B73. (Anderson et al. 2021). To categorize gene expression categories used in Figure 5C, a B73 developmental expression atlas data (Stelpflug et al. 2016) was analyzed using similar methods as previously (Anderson et al. 2019b). Genes that are constitutively lowly or not expressed (“constitutive off”) have an RPM > 1 in fewer than 3 of the 225 total libraries representing different tissues, developmental stages, and biological replicates. Endosperm expressed (endosperm on) and not endosperm expressed (endosperm off) were identified from the rest of the genes based on the following criteria: The “endosperm on” genes have an RPM > 1 in at least 18 of 21 endosperm libraries. The “endosperm off” genes have an RPM value of < 0.5 in at least 20 of the 21 endosperm libraries. Finally, the W22 genes corresponding to the B73 genes meeting these criteria were determined with a gene key, which can be found at https://github.com/SNAnderson/Expression/blob/main/genes.expression.development.categories.20Sept21.txt

### Methylome read processing, definition of regions by methylation status

EM-seq reads were trimmed of adapter sequence using cutadapt, parameters -q 20 -a AGATCGGAAGAGC -A AGATCGGAAGAGC -O. Reads were aligned to each genome and methylation values called using BS-Seeker2 (version 2.1.5), parameters -m 1 --aligner=bowtie2 - X 1000 (Guo et al. 2013). The quality of the enzymatic conversion for each biological replicate was validated using mCHH values from the unmenthylated lambda spike-in control (Supplemental Table 3) and by examining methylation near gene transcription start sites as in Supplemental Figure S3A. To identify DMRs, CGmaps from each group of three individual replicates were merged into single CGmaps using the merge2 tool of CGmapTools version 0.1.2 (Guo et al. 2018). Intersected CGmaps for each pair of merged CGmaps were produced using the CGmapTools intersect tool in each sequence context separately (CG, CHG, and CHH). Single-bp level methylation measurements in lollipop plots were produced using the CGmapTools lollipop tool on intersected CGmaps. CG and CHG DMRs and control eligible regions were then identified from the separate CG and CHG intersected maps using DMR_Finder.py, parameters as follows: binSize = 200, minCov = 3 minCountC = 5, minAbsDif = .20, minRelDif = .5. The set of CG DMRs that overlapped CHG DMRs were identified using the BEDTools (version 2.29.2) intersect tool. To finalize the set of DMRs, all DMRs located on scaffolds rather than chromosomes 1 through 10 were discarded. mCHHregions were identified from the merged endosperm CGmap using MR_Finder.py (see Supplemental scripts), parameters as follows: binSize = 200, minCov = 2, minCountC = 5, minMeth = .2, context1 = ‘CHH’, context2 = ‘CHH’. CHG unmethylated regions (UMRs) were identified from the merged embryo CGmap using UMR_Finder.py (see Supplemental scripts), parameters as follows: binSize = 200, minCov = 2, minCountC = 5, maxMeth = .2, context1 = ‘CHG’, context2 = ‘CHG’. Scripts are available at https://github.com/dawelab/mdr1.

### Characterization of DMRs in terms of methylation levels, overlap with specific genetic elements, and sequence motifs

Methylation averages on 100-bp intervals centered on DMRs and control eligible regions were produced using the CGmapTools mfg tool, requiring a minimum coverage of one with the -c 1 parameter. The numbers of overlaps between regions and different genetic elements, requiring at least 100 bp of overlap to count, were obtained using the BEDTools intersect tool with parameters -u -f .5. To determine whether DMRs contained motifs associated with differential methylation, fasta sequences were derived from bed files using the BEDTools getfasta tool.

Three subsamples of 5000 sequences were input into STREME, each with a different seed number (Bailey 2021). All motifs identified by STREME were input into FIMO to identify matches throughout the complete sets of sequences. SpaMo (Whitington et al. 2011) was then used to search for combinations of motifs that may work together to cause the differential methylation, but was also inconclusive. AT richness was measured using the EMBOSS program infoseq (Rice et al. 2000).

### Characterization of siRNA expression in methylated regions

Small RNA-seq reads were quality filtered, trimmed of adapters, and filtered for lengths of 20-25 nt using cutadapt (Martin, 2011), parameters -q 20 -a TGGAATTCTCGGGTGCCAAGG -e .05 -O 5 --discard-untrimmed -m 28 -M 33. The four random nucleotides at each end were then removed using cutadapt with the -u 4 parameter followed by cutadapt with the -u −4 parameter. Reads were aligned to the genome with Bowtie2, --very-sensitive parameters. Reads that overlapped at least 90% of their lengths with tRNA, 5S RNA, NOR, or miRNA loci were removed using the BEDTools intersect tool with parameters -v -f .9. The remaining reads were called siRNA reads. Reads that overlapped DMRs were identified the same way. For the analysis of uniquely mapping siRNAs reads overlapping DMRs, reads with a MAPQ score of q20 were selected.

## Accession numbers

All raw sequencing data generated in this study have been submitted to the NCBI BioProject database (https://www.ncbi.nlm.nih.gov/bioproject/) under accession number PRJNA759188. Individual accession numbers for DNA methylation are listed in Supplemental Table S3 and for small RNAs in Supplemental Table S4.

## ACKNOWLEDGMENTS

We thank the Maize Genetics Cooperation Stock Center for providing maize stocks; Akshay Nair, Ankush Sangra, Olivia Smith, and Mary Tindall Smith for genotyping stocks; Justin Scherer for help with making protein trees; and John Fowler for advice on use of Ds-GFP insertions for measuring frequency of mutation transmission. This study was supported in part by resources and technical expertise from the Georgia Advanced Computing Resource Center, a partnership between the University of Georgia’s Office of the Vice President for Research and Office of the Vice President for Information Technology.

