## Supplemental Figures 1-8 for "The maize gene *maternal derepression of r1* (*mdr1*) encodes a DNA glycosylase that demethylates DNA and reduces siRNA expression in endosperm"

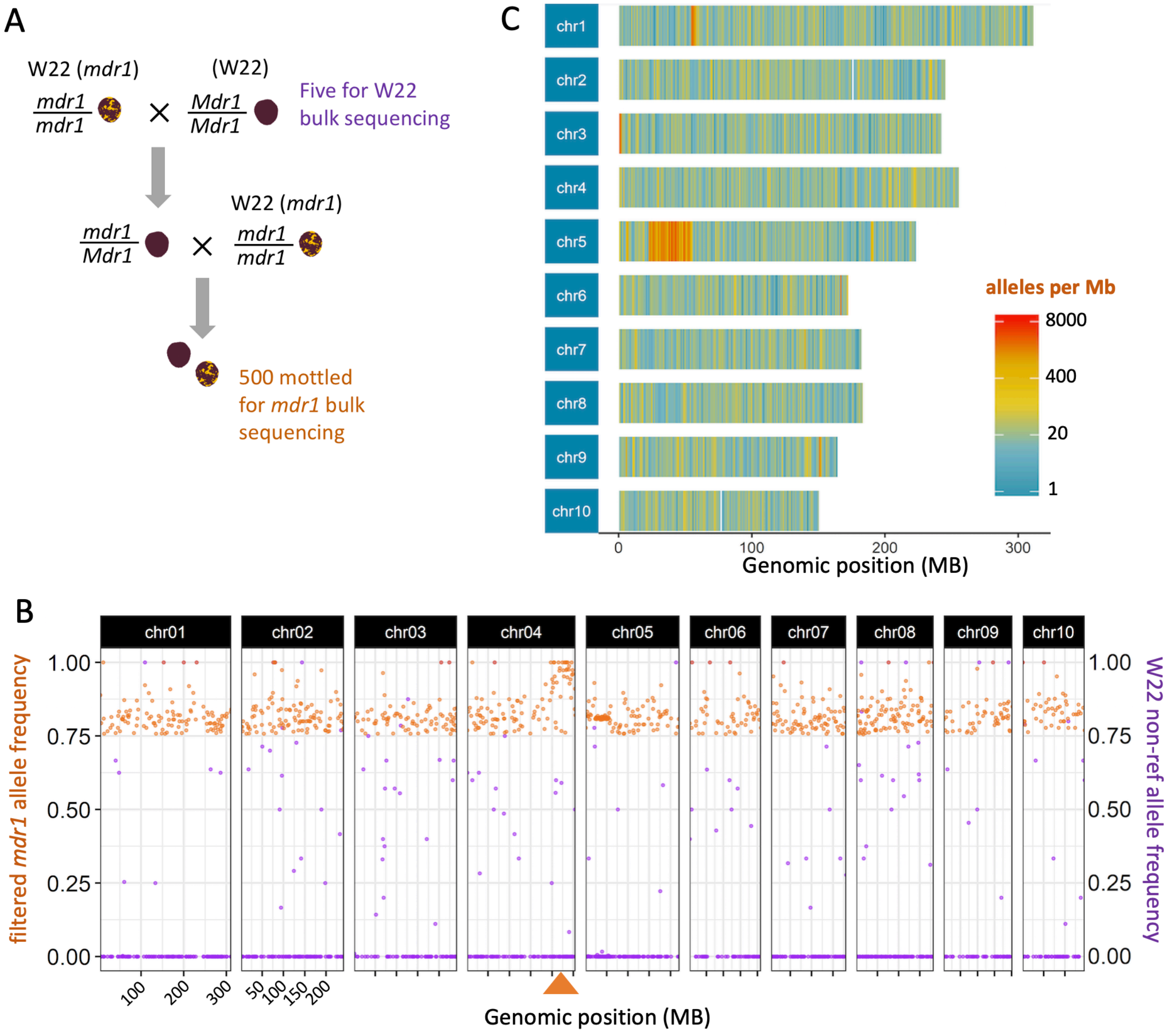


**Figure S1**

**A,** Schematic depiction of *mdr1* mapping method based on kernel mottling phenotype.

**B,** Allele frequency is the number of reads supporting the non-W22-reference allele divided by the total number of reads at that site, averaged over 1-Mb intervals. The left Y-axis (orange) indicates variant allele frequencies for the *mdr1* bulk, filtered to only include ones with a frequency greater than 0.75. Values are averaged over non-overlapping 1 Mb intervals. According to the crossing scheme (see panel A), *mdr1*-linked alleles should approach frequencies of 1, while unlinked ones should average 0.75. The right Y-axis (purple) indicates allele frequency for the W22 bulk for all variant alelles, not just ones with a frequency greater than 0.75. The X-axis indicates genomic position, with major tick marks on 50-Mb intervals, except for chromosome 1 on 100-Mb intervals. The arrowhead on the distal tip of chromosome 4 indicates position of *dng101*/*mdr1*.

**C,** Chromosome heatmaps indicate the number of non-W22-reference alleles per Mb in the *mdr1* bulk. High frequency alleles reveal regions that are not shared between the sequenced W22 stock and Kermicles's *mdr1* W22 stock. These regions do not include the *mdr1* region on the tip of 4L.

| maternal genotype | | paternal genotype | viable progeny that inherited both mutations | |
|---|---|---|---|---|
| *Mdr1*/*Mdr1* *Dng102*/*Dng102* | X | *Mdr1*/*mdr1* *Dng102*/*mu1083641* | 0 of 35 | (P-value = 0.0003) |
| *Mdr1*/*mdr1* *Dng102*/*mu1083641* | X | *Mdr1*/*Mdr1* *Dng102*/*Dng102* | 0 of 72 | (P-value < 0.0001) |
| *Mdr1*/*Mdr1* *Dng102*/*Dng102* | X | *Mdr1*/*mdr1* *Dng102*/*dng102-Q235* | 0 of 23 | (P-value = 0.0028) |
| *Mdr1*/*mdr1* *Dng102*/*dng102-Q235* | X | *Mdr1*/*Mdr1* *Dng102*/*Dng102* | 0 of 46 | (P-value < 0.0001) |

**Figure S2**
Crosses between heterozygous double mutant *mdr1 dng102* and wild-type fail to transmit both mutations simultaneously. Kernels from each cross were planted and seedling leaves genotyped by PCR. P-values are based on a Fisher's exact test compared to the expected frequency of 25% double mutant transmission.

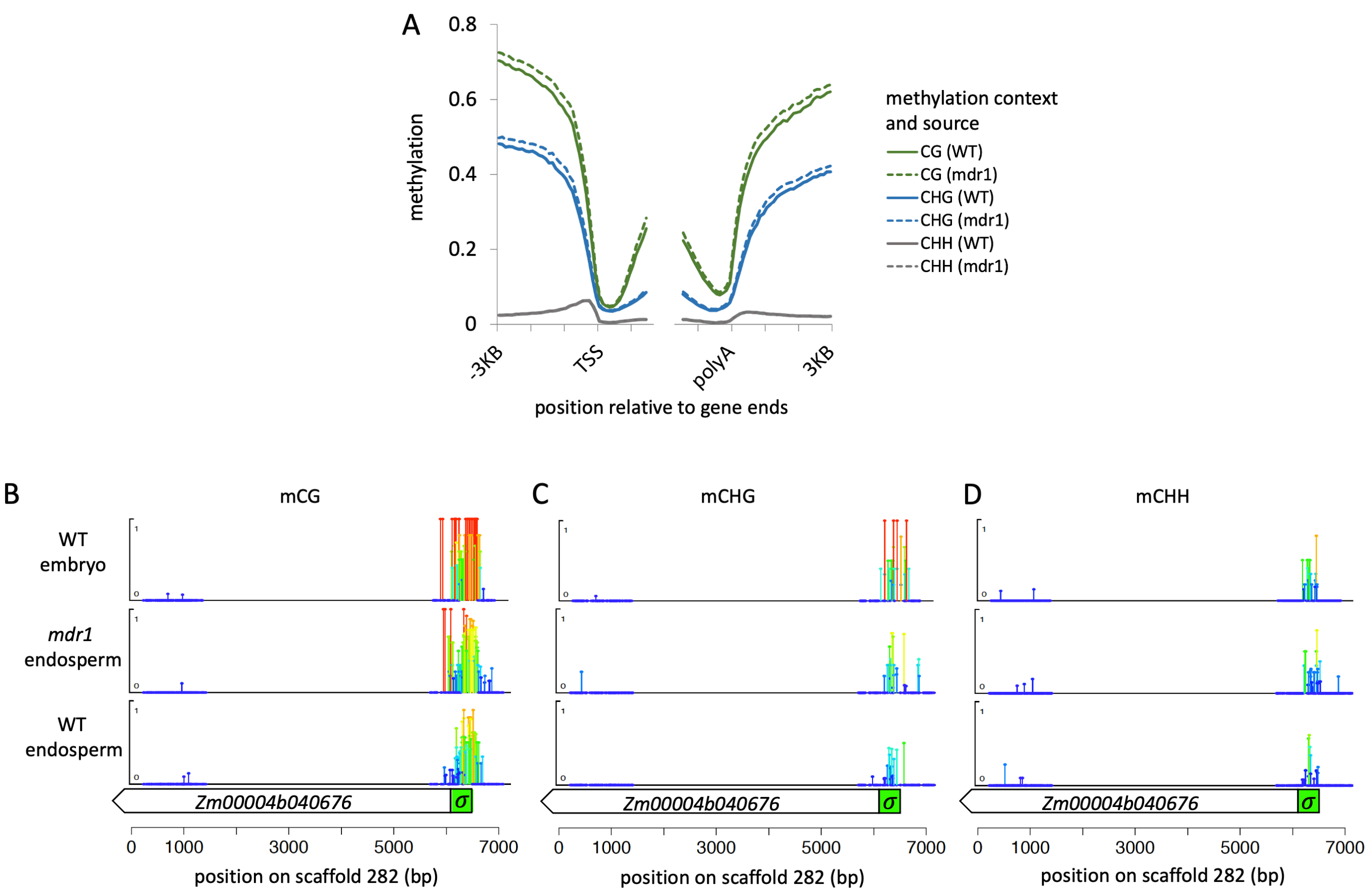


**Figure S3**

**A,** Wild-type and *mdr1* mutant mCG, mCHG, and mCHH profiles relative to gene ends. Methylation values are averages of 100-bp intervals. TSS = Transcription start site, polyA = polyadenylation site.

**B**, Differential methylation of an *r1* gene in embryo, endosperm, and *mdr1* mutant endosperm, same as Figure 3A. mCG values are shown at single-base level resolution over an 8-Kb region including the majority of an *r1* homolog in W22 (*Zm00004b040676*). Both height and color of lollipops indicate mCG values (darkest blue = zero mCG). Regions without lollipops indicate lack of read coverage, either because of multi-copy sequence or large N gaps.

**C**, Same as B, but mCHG.

**D,** Same as A, but mCHH.

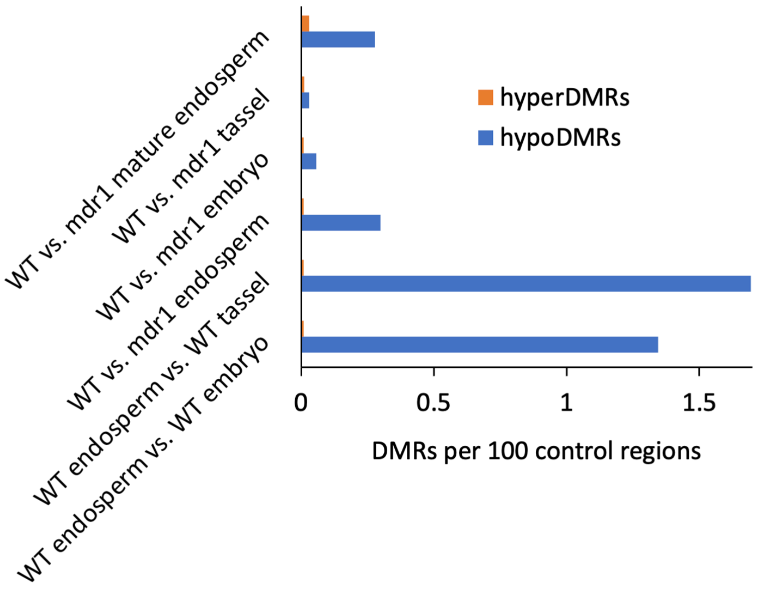


**Figure S4**

HypoDMR and hyperDMR frequencies for pairwise comparisons of multiple tissues and genotypes. Control regions are the 200-bp regions that were eligible for identifying DMRs based on read coverage and number of informative cytosines. Mature endosperm is from dry seeds. Other endosperm samples are 15-DAP. Embryos are also 15-DAP. Tassel is premeiotic stage.

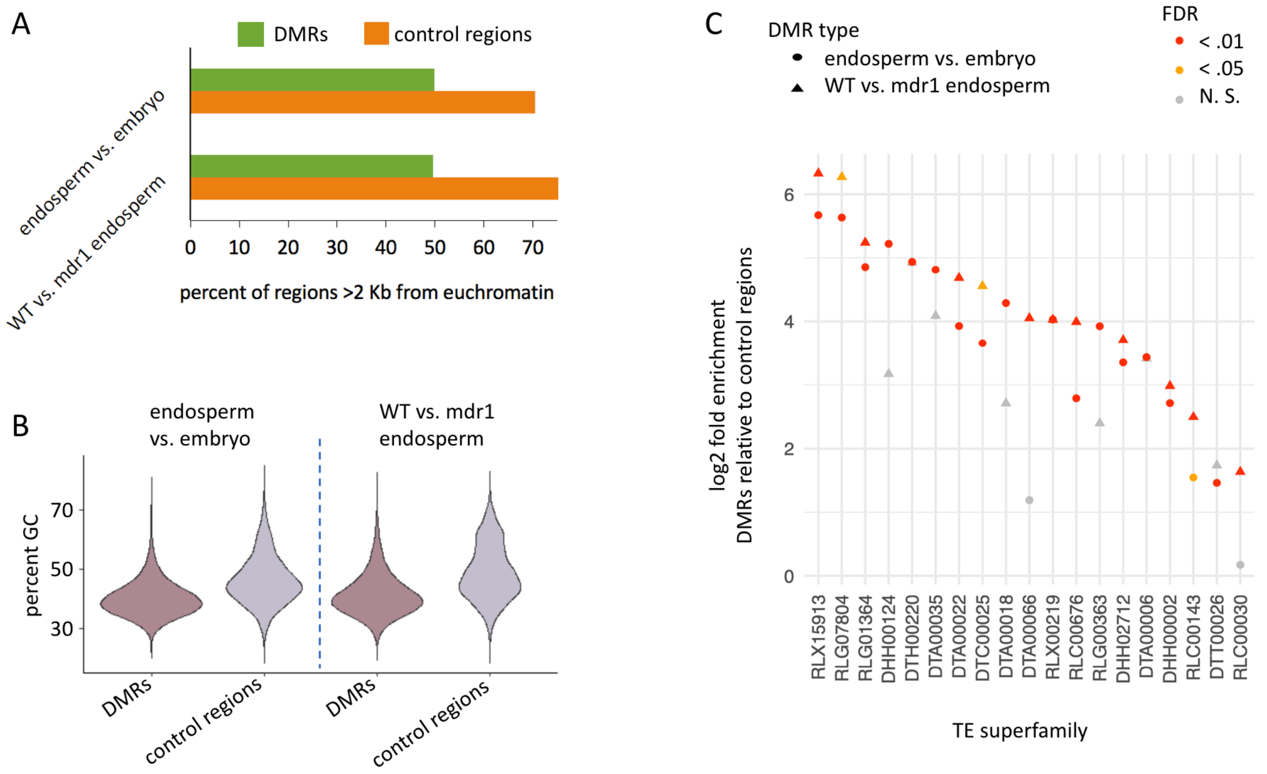


**Figure S5**

**A**, Percent of regions that are greater than 2 Kb from nearest euchromatin. Euchromatin is defined by 200 bp regions with mCHG < 0.2 in 15-DAP embryo.

**B**, GC content of DMRs and control regions.

**C**, DMR enrichment in different transposon families. Enrichment is the proportion of DMRs that overlap at least 50% of their length (100 bp) with elements of the specified families divided by the proportion of control regions that do. Control regions are the 200-bp regions that were eligible for identifying DMRs based on read coverage and number of informative cytosines. Three letter abbreviations in family names indicate superfamilies: DTA is hAT, DTC is CACTA, DTT is Tc1/Mariner, RLC is Copia, RLG is Gypsy, and RLX is unclassified LTR retrotransposon. Only families with at least three copies that overlap DMRs and have at least 10 DMRs total in at least one comparison are included. Some copies overlap more than one DMR. P-values were calculated with the binomial test.

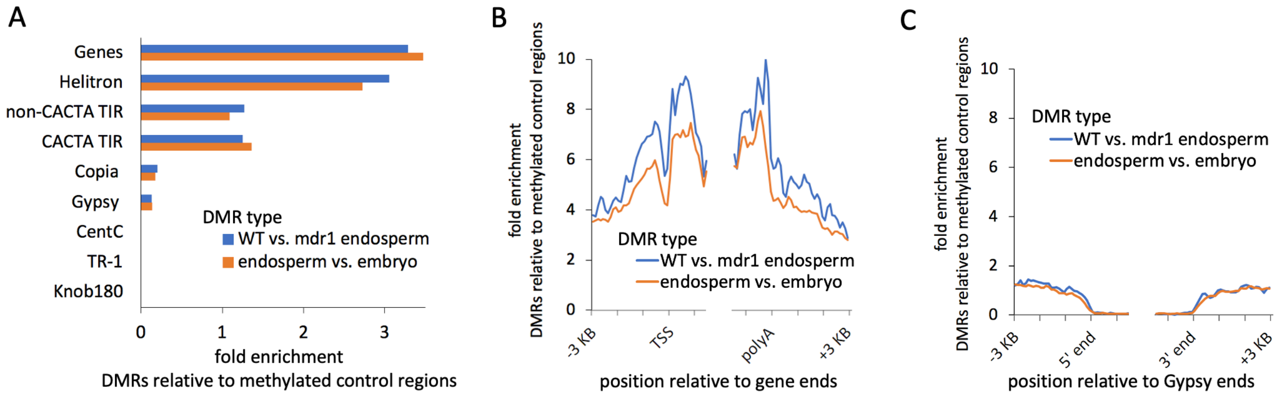


**Figure S6**

**A**, DMR enrichment relative to methylated control regions in different genetic elements. Enrichment is the proportion of DMRs that overlap at least 50% of their length (100 bp) divided by the proportion of methylated control regions that do. Methylated control regions are the 200-bp eligible regions used to define DMRs, but only including the subset where both mCG and mCHG exceeded values of 0.2 in at least one of the two methylomes. **B,C,** Enrichment for DMRs relative to methylated control regions on 100-bp intervals relative to gene ends (C) or Gypsy ends (D). Gypsies with solo LTRs were excluded from this analysis. For genes, TSS = transcription start site, polyA = polyadenylation site.

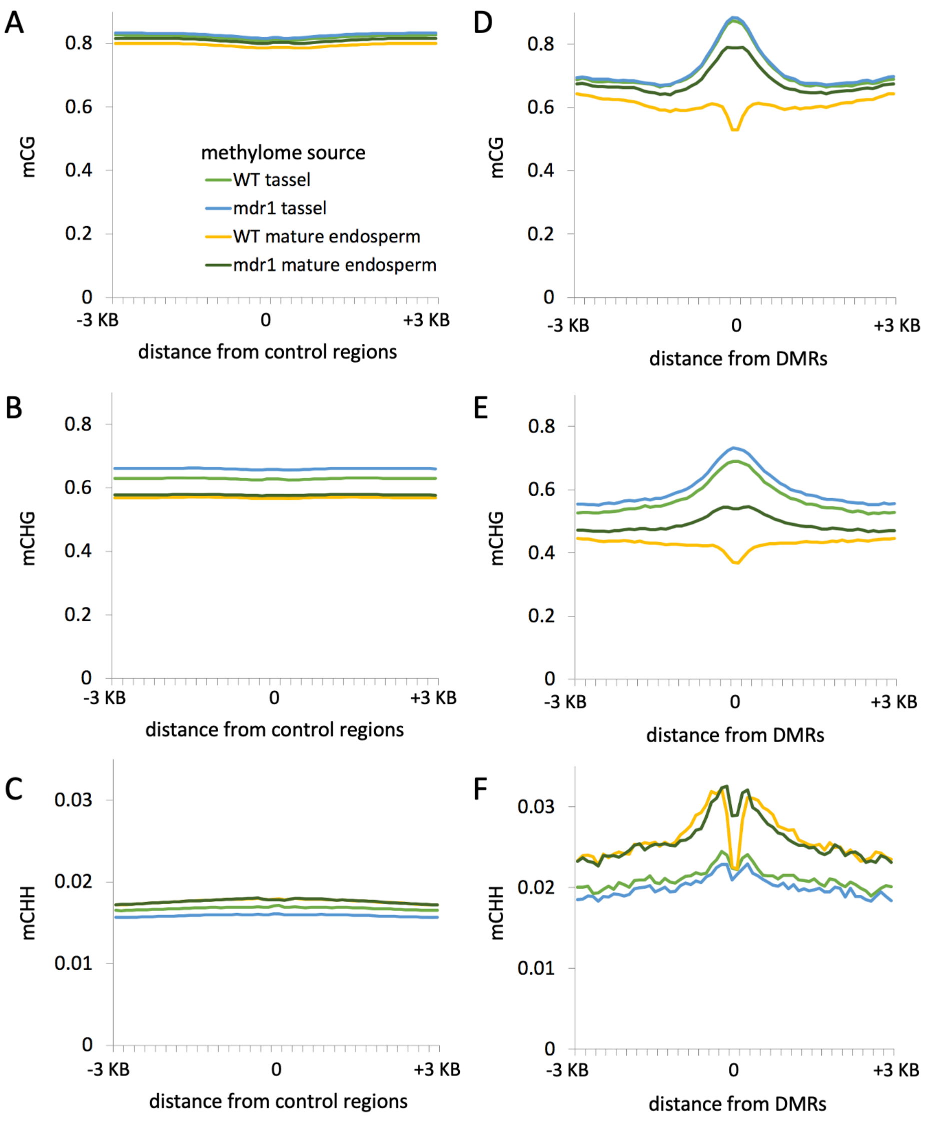


**Figure S7**

**A-C,** mCG, mCHG, and mCHH profiles centered on control regions associated with wild-type vs. mdr1 endosperm DMRs. Methylation values are averages of 100-bp intervals within and up to 3 Kb on either side of each 200-bp region. Mature endosperm is from dry seeds. Tassel is premeiotic stage.

**D-F,** Same as A-C, except methylation centered on wild-type vs. mdr1 DMRs instead of control regions.

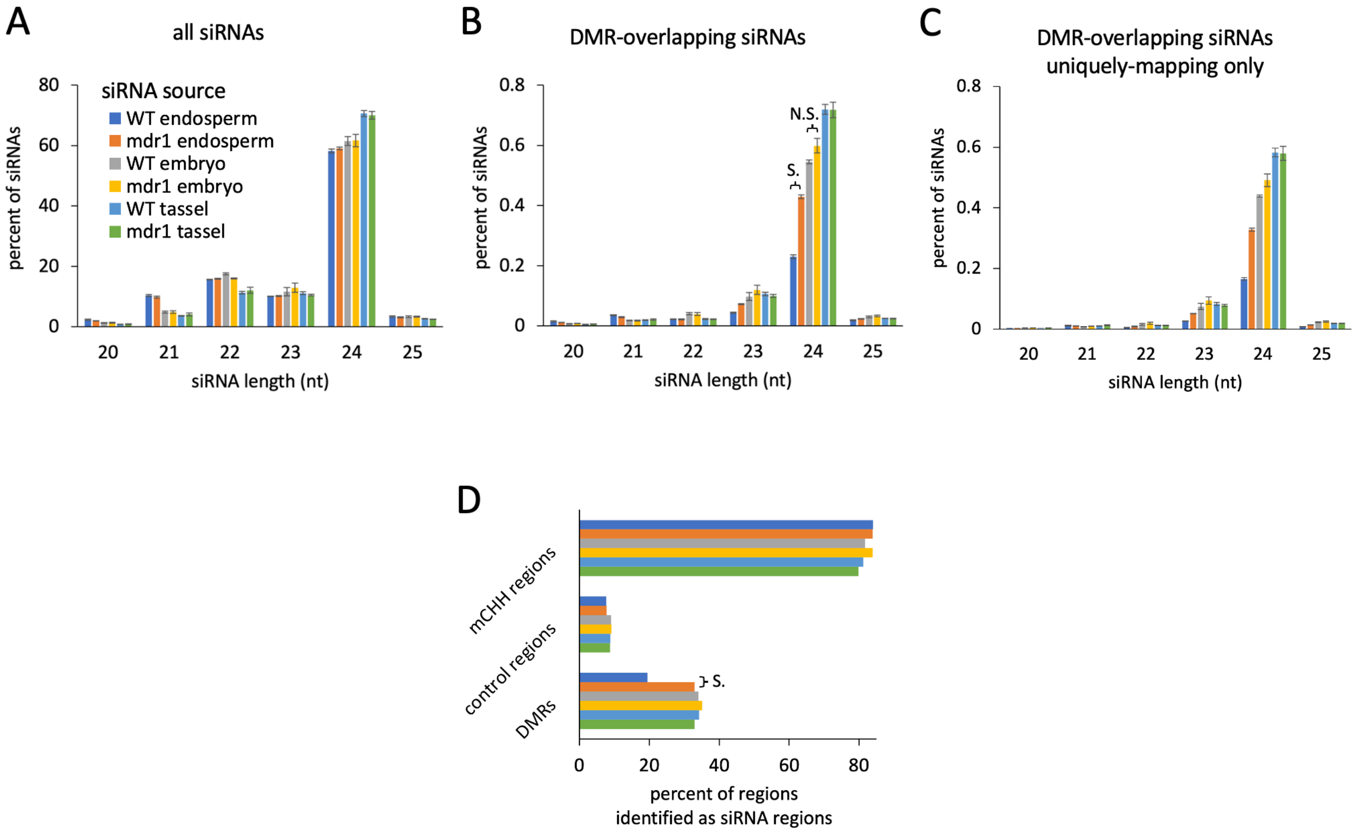


**Figure S8**

**A**, Length distributions of all siRNAs. Error bars are standard errors of the means for biological replicates. Same as Figure 4G, except including premeiotic stage tassel siRNAs.

**B**, As in A, except only siRNAs that overlapped at least 90% of their length with wild-type vs. mdr1 endosperm DMRs. “S.” indicates significant difference: P-value < .00001, two-tailed Student t-test. “N.S.” indicated not significant: P-value = .06. Same as Figure 4H, except including premeiotic stage tassel siRNAs.

**C**, As in B, but only including uniquely-mapping siRNAs.

**D**, Regions categorized by whether they produce siRNAs. An “siRNA region” is any that had at least 50 bp spanned by siRNAs of the indicated source tissues. All libraries were subsampled to 20 million siRNA reads. DMRs and control regions are from the wild-type vs. mdr1 endosperm comparison. mCHH regions are 200-bp regions with an average mCHH value of at least 0.2 in wild-type endosperm. Same as Figure 4I, except including premeiotic stage tassel siRNAs. “S.” indicates significant difference (P-value < .0001 two-tailed Chi-Square with Yates correction).
